# GATHeR: Graph-based Accurate Tool for Immunoglobulin HEavy- and Light-chain Reconstruction

**DOI:** 10.1101/2025.09.21.677530

**Authors:** Seyedmojtaba Seyedraoufi, Mari Bergstøl Gornitzka, Andreas Lossius

## Abstract

Recovering full-length, paired B-cell receptor (BCR) sequences from scRNA-seq reads remains difficult, especially in naive and memory B cells where immunoglobulin transcripts are sparse. Current methods provide incomplete constant-region coverage, limiting isoform, subclass, and allele resolution. We present GATHeR, an open-source tool that assembles and annotates paired heavy- and light-chain BCR sequences and extends assembled sequences into constant regions. This enables confident subclass and allele assignment and recovery of the membrane-bound isoform, including the transmembrane segment and cytoplasmic tail, thereby distinguishing surface BCRs from secreted antibodies. GATHeR supports Smart-seq2/3 and 10x Genomics libraries and outperforms existing methods across benchmarks, with the largest gains in naive and memory B cells. Notably, in these populations the constant-region extension also revealed splice variation, including heavy-chain intron retention. By delivering high-fidelity receptor, isoform, and clonal lineage information, GATHeR broadens the analytical reach of scRNA-seq for B-cell immunology.

## Introduction

Humoral immunity depends on antigen recognition by the B-cell receptor (BCR), which comprises a membrane immunoglobulin (Ig)—a heterotetramer of two identical heavy and two identical light chains with variable (V) and constant (C) regions—associated with the CD79a/CD79b (Ig*α*/Ig*β*) signaling heterodimer. Following BCR engagement by cognate antigen and receipt of T cell help, naive B cells either generate short-lived extrafollicular plasmablasts or seed germinal centers for affinity maturation and selection, ultimately producing long-lived plasma cells and memory B cells [1]. Commitment to antibody-secreting fates entails extensive remodeling of protein synthesis and secretory pathways. Accordingly, Ig transcripts can account for up to 70% of total mRNA in antibody-secreting cells versus ~ 2% in naive or memory populations [2]. This transition is accompanied by regulated alternative splicing and polyadenylation of Ig heavy-chain pre-mRNA at the 3′ end, which selects the secretory tailpiece rather than the membrane exons (M1/M2) and thus produces the secreted Ig isoform [3].

Single-cell RNA sequencing (scRNA-seq) now enables joint profiling of cell states and receptor usage. For B cells, accurately recovering paired heavy- and light-chain sequences together with the transcriptome is essential for functional annotation and clonal tracing. To achieve this, standard droplet-based workflows such as 10x Genomics require researchers to prepare additional receptor-specific libraries to capture BCR sequences [4]. Because these assays amplify from CH1-anchored primers, determining Ig subclasses and alleles is difficult—often impossible. Meanwhile, several full-length scRNA-seq protocols (e.g., Smart-seq2/3 [5, 6] and FLASH-seq [7]) have been developed, but in these datasets, BCR transcripts are effectively ‘buried’ within the complete transcriptome and must be extracted and correctly assembled.

Here we present GATHeR, an open-source pipeline that reconstructs paired, full-length BCRs directly from scRNA-seq. GATHeR natively supports full-length protocols (Smart-seq2/3) and also assembles BCRs from 10x Genomics 5′ gene expression libraries without additional receptor-specific enrichment. Across diverse datasets and compared with BASIC [8], BraCeR [9], and TRUST4 [10], GATHeR yields markedly longer, more contiguous BCR assemblies and higher assembly success rates, with the largest gains in naive and memory B cells where Ig transcripts are sparse. Crucially, when constant-region reads are available, the assemblies extend across the constant region, enabling interrogation of constant-region isoforms, subclasses and alleles. In our analyses, these long assemblies also revealed previously unreported splice variation in naive and memory B cells, including heavy-chain intron retention.

## Results

### GATHeR assembles and annotates full-length BCRs and supports phylogenetic analysis

GATHeR starts by building a compacted de Bruijn graph (cDBG) from raw scRNA-seq reads (Figure 1A). Traversing cDBG subgraphs enables recovery of contigs that contain complete Ig transcripts, including the transmembrane and cytoplasmic-tail exons of membrane-bound IgM and IgD in naive B cells. Candidate heavy- and light-chain contigs are identified by pairwise alignment to the IMGT germline database, annotated with V(D)J gene usage, constant-region subclass/allele, and average contig weight (derived from k-mer frequencies), which is recorded in the FASTA header; secondary contigs are retained whenever multiple heavy or light chains are detected.

**Figure 1.**
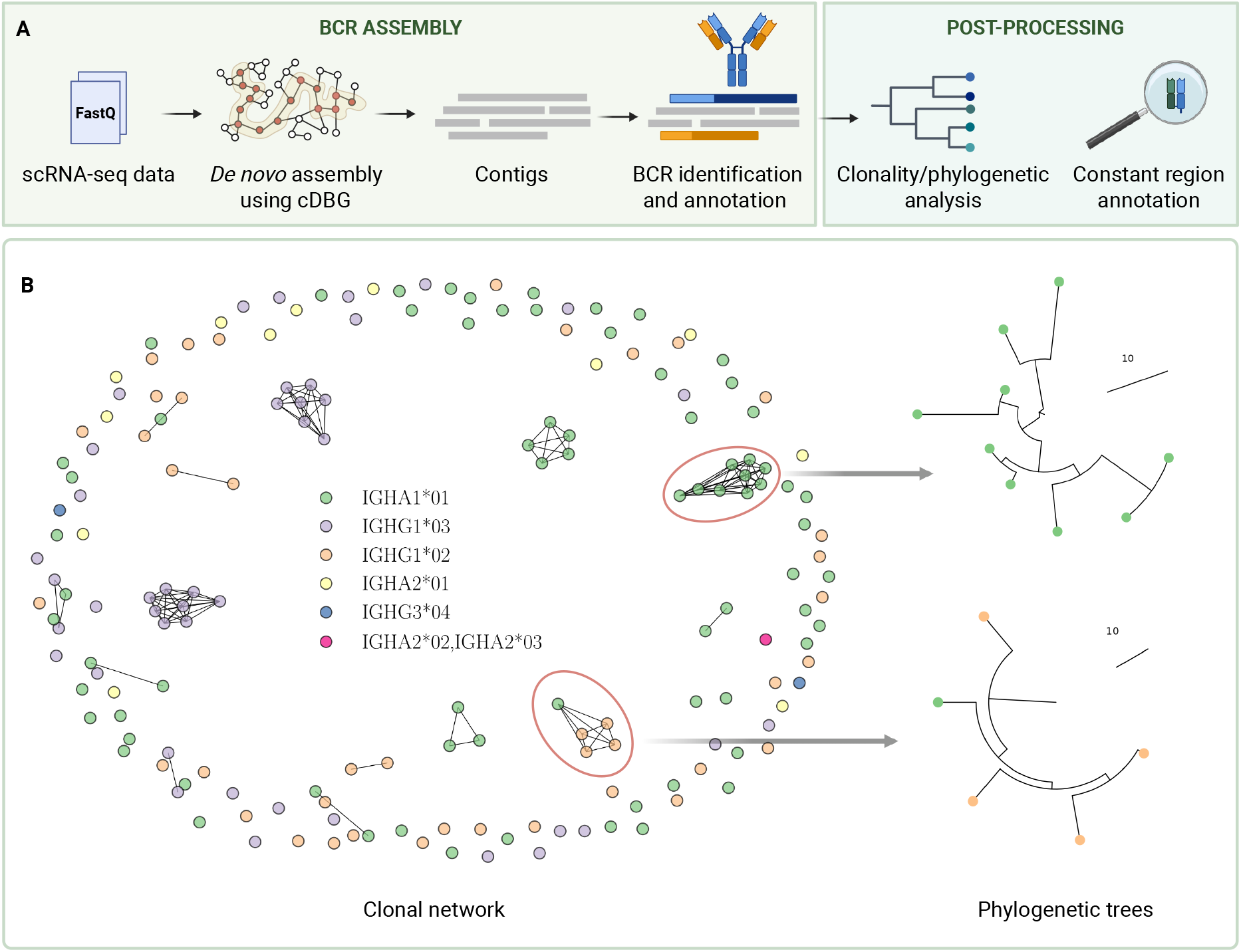
GATHeR reconstructs paired full-length BCRs from scRNA-seq and enables clonality and lineage analysis. (**A**) Overview of the GATHeR workflow. (**B**) Clonal network of B cells derived from Ig clonality analysis of dataset I in Table 1 (174 human plasmablasts) [8]. Nodes are colored according to Ig isotype. Phylogenetic trees for two highlighted clones are shown on the right.

A post-processing module wraps Immcantation tools to cluster BCR contigs into clonal families and reconstruct maximum-parsimony lineage trees (Figure 1B), providing a seamless path from raw reads to repertoire-scale phylogenetic analysis. Importantly, GATHeR annotates all constant-region exons against the IMGT reference, and is able to infer novel constant region alleles (Supplementary Table 1).

### GATHeR achieves high reconstruction yield and sequence-level concordance with existing methods

To evaluate GATHeR, we analyzed three publicly available full-length scRNA-seq datasets and one 10x Genomics 5′ gene expression dataset (Datasets I–IV; Table 1). We benchmarked against three established BCR assembly tools selected based on favorable performance in a recent comparative study [11]: BASIC [8], BraCeR [9], and TRUST4 [10]. Dataset I is the original validation set for BASIC [8], and Dataset II corresponds to the data used in the BraCeR publication [9]. Because both datasets consist exclusively of antibody-secreting cells and thus have high Ig transcript abundance, we also included Dataset III, randomly sampling 200 naive and 200 memory B cells (1:1) to assess performance in settings with sparse Ig transcripts. Finally, to assess compatibility with 10x Genomics 5′ gene-expression libraries lacking receptor-specific enrichment, we included Dataset IV, which was previously used to validate TRUST4 [10].

**Table 1.** Datasets used to evaluate the performance of GATHeR.

| Dataset | Platform | Species | Cell type(s) | Cell count | Reference |
| --- | --- | --- | --- | --- | --- |
| I | Smart-seq2 | Human | Plasmablasts | 174 | Canzar et al. [8] |
| II | Smart-seq2 | Mouse | IL4/LPS stimulated B cells | 56 | Wu et al. [12] |
| III | Smart-seq3xpress | Human | naive and memory B cells (1:1) | 400 | Hagemann-Jensen et al. [6] |
| IV | 10× Genomics 5' | Human | naive and memory B cells, plasmablasts | 1,111 | 10x Genomics [13] |

Across Datasets I–II, all tools reconstructed nearly all paired heavy–light chains (Supplementary Figure 1A). In Dataset III (naive and memory B cells), GATHeR assembled BCRs for 399/400 cells (one heavy missing) and TRUST4 for 398/400 (two heavy missing); BASIC assembled 94% (23 heavy, 24 light missing), while BraCeR assembled 78% heavy (312/400) and 96% light (383/400). Sequence-level concordance with GATHeR was high across all datasets (Supplementary Figure 1B,C): BraCeR had 98–100% of contigs with similarity *≥* 0.9, TRUST4 had 93–98% (heavy) and 91–98% (light), and BASIC was lower in Datasets I–II (64–84% heavy; 66–79% light) but ~ 99.7% in Dataset III. For cells with multiple or fragmented contigs (most notably with BASIC), we compared all contigs to the GATHeR assembly and reported the maximum per cell.

### GATHeR reconstructs longer, more complete BCRs, including membrane-bound IgM/IgD isoforms

Long, contiguous BCR sequences that span the variable region and the entire constant region of both heavy and light chains enable confident isotype/subclass and allele assignment and support robust clonal lineage analysis. Across Datasets I–III, GATHeR typically produced full-length heavy- and light-chain sequences (median normalized length *≈* 1.0; Figure 2A). BASIC approached full length in Dataset II, whereas BraCeR and TRUST4 yielded shorter sequences across datasets. Consistently, heavy-chain constant-region coverage (Figure 2B) mirrored these differences, with GATHeR recovering the longest segments across datasets.

**Figure 2.**
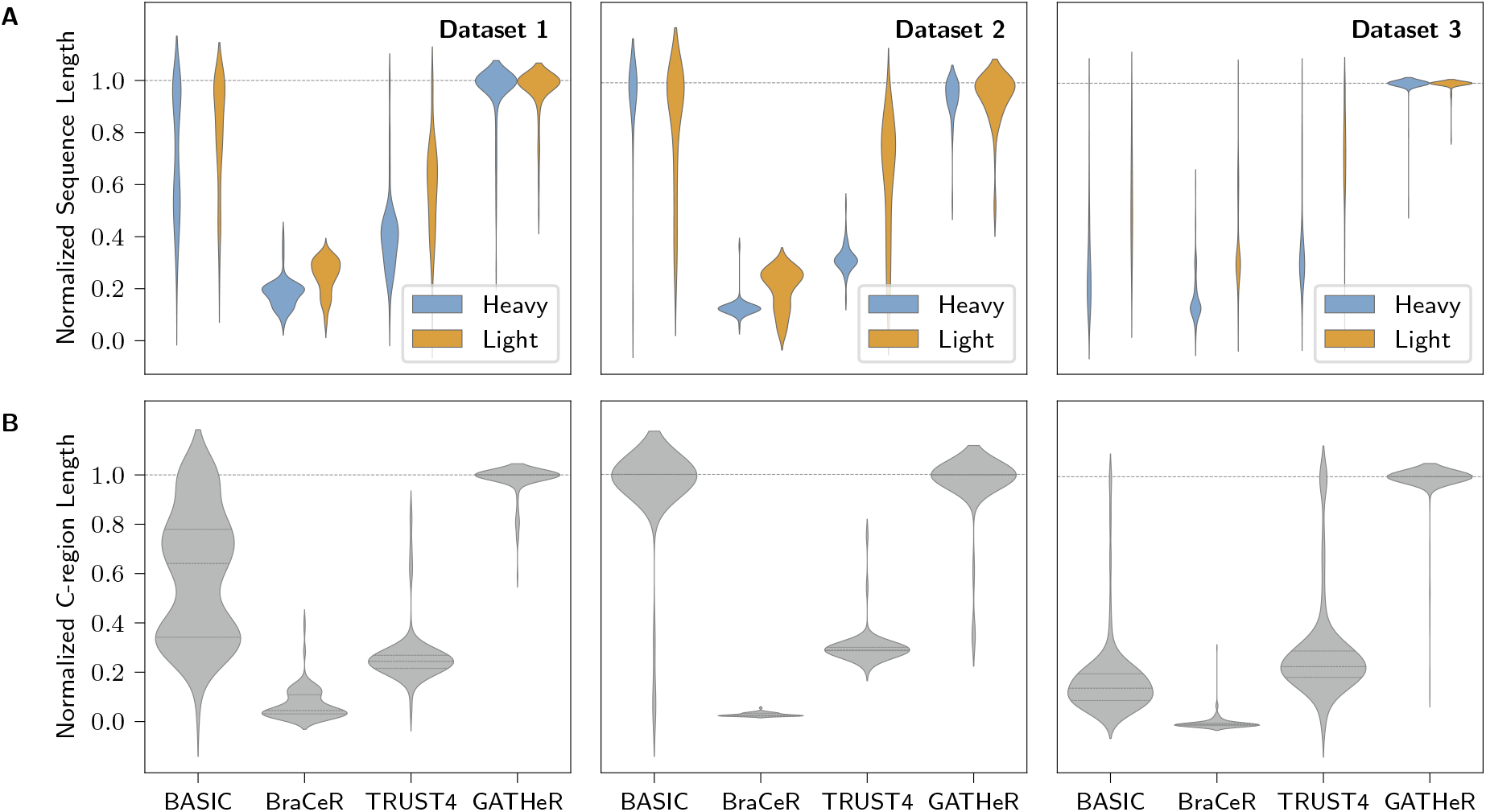
GATHeR reconstructs longer, more complete heavy- and light chains across datasets. (**A**) Distributions of contig lengths for heavy- and light-chain assemblies produced by GATHeR and benchmark methods (BASIC, BraCeR, TRUST4) across Datasets I–III. For each cell and chain type, lengths are normalized to the longest contig produced by any method for that cell (so 1 denotes the longest per-cell contig among methods). (**B**) Corresponding heavy-chain constant-region coverage, shown as normalized length distributions using the same per-cell normalization.

Sequence integrity differed markedly among methods in the naive and memory B cells in Dataset III (Figure 3A). GATHeR delivered the highest contiguity, assembling heavy- and light-chain sequences as contiguous in 93% and 100% of cells, respectively. By contrast, BASIC achieved contiguity in only 9% (heavy chains) and 35% (light chains) of cells; TRUST4 closely matched GATHeR, with marginally lower contiguity for heavy chains, while BraCeR produced mostly contiguous heavy and light chains but at a lower overall success rate.

**Figure 3.**
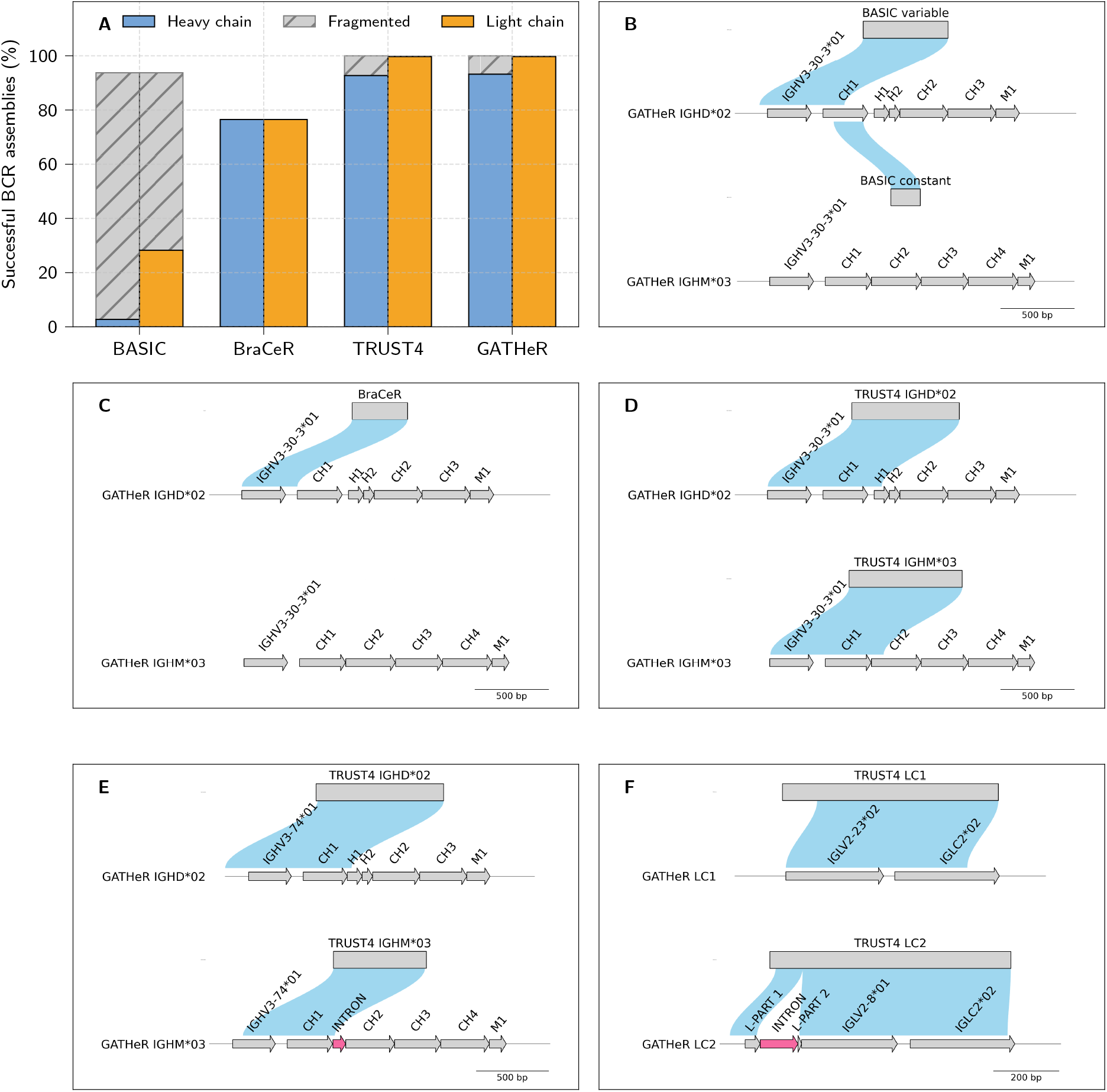
GATHeR preserves sequence contiguity and resolves intron retention in BCR assemblies. (**A**) Percentage of cells with contiguous heavy- and light-chain assemblies for each method in Dataset III (naive and memory B cells); the fraction of fragmented assemblies is indicated in gray. (**B–D**) Representative naive B cell: GATHeR-assembled IgM and IgD heavy-chain sequences with assemblies from other methods aligned above: (**B**) fragmented BASIC, (**C**) BraCeR, and (**D**) TRUST4. (**E**) Example of intron retention in the IgM constant region detected by GATHeR; the aligned TRUST4 assemblies above miss the retained intron. (**F**) Light-chain example with a partial retained intron between L-PART1 and L-PART2 detected by GATHeR; the aligned TRUST4 assembly lacks this event.

Previous studies have identified co-expression of multiple BCRs in a subset of mature naive and memory B cells [14, 15]. In Dataset III, GATHeR identified two or more heavy chains in 35% of memory and 55% of naive B cells (Supplementary Figure 2A), and two or more light chains in 33% and 24%, respectively (Supplementary Figure 2B). Among naive B cells with multiple heavy chains, most cases comprised distinct IGHM and IGHD transcripts sharing an identical VDJ rearrangement, consistent with IgM/IgD co-expression in mature naive B cells [16]. Panels B–D of Figure 3 illustrate BCR reconstruction in a representative naive B cell: GATHeR reconstructs two full-length heavy-chain isoforms (IgM and IgD), whereas BASIC (Figure 3B) splits one into variable- and constant-region fragments, and BraCeR (Figure 3C) and TRUST4 (Figure 3D) return shorter single-contig reconstructions.

GATHeR’s extended heavy- and light-chain sequences also enabled us to resolve alternative splicing. In Dataset III, intron retention was detected in heavy-chain sequences in 8 % of naive and 9 % of memory B cells (Supplementary Figure 3), and in light-chain sequences in 10 % and 23 % of cells, respectively (Supplementary Figure 4). Among intron-retained sequences, 37/38 heavy chains (97%) and 77/79 light chains (97%) harbored a frameshift and/or a premature termination codon, thereby contributing to non-productive transcripts in these B-cell populations (Supplementary Figure 2C,D). Figure 3E illustrates intron retention in the constant region of a GATHeR-reconstructed IgM from a naive B cell; the corresponding TRUST4 reconstruction missed this event, potentially yielding a false productive call. A representative light-chain example is shown in Figure 3F: GATHeR reconstructs a light chain with a partial intron between the first and second leader exons (L-PART1 and L-PART2) that TRUST4 does not capture; the partial intron introduces a premature termination codon, making the transcript non-productive. These observations align with a prior report of light-chain intron retention in naive B cells [17] and extend it to heavy chains in both naive and memory B cells.

### GATHeR successfully assembles BCRs from 10x Genomics 5^′^ gene expression libraries

Unlike the standard 10x Genomics workflow, which typically relies on an additional V(D)J-enriched library, GATHeR reconstructs BCRs directly from 5′ gene-expression (GEX) data, eliminating that step. Because TRUST4 is the only comparator with native GEX support, we benchmarked GATHeR against TRUST4 on 1,111 Seurat-annotated naive and memory B cells together with a small proportion of plasmablasts (Figure 4A; Table 1, Dataset IV). Both methods assembled at least one chain in *>* 99% of cells (Figure 4B). For sequences spanning CDR3, GATHeR reconstructed heavy and light chains in 70.5% and 99.0% of cells, respectively (TRUST4: 70.1% and 99.5%).

**Figure 4.**
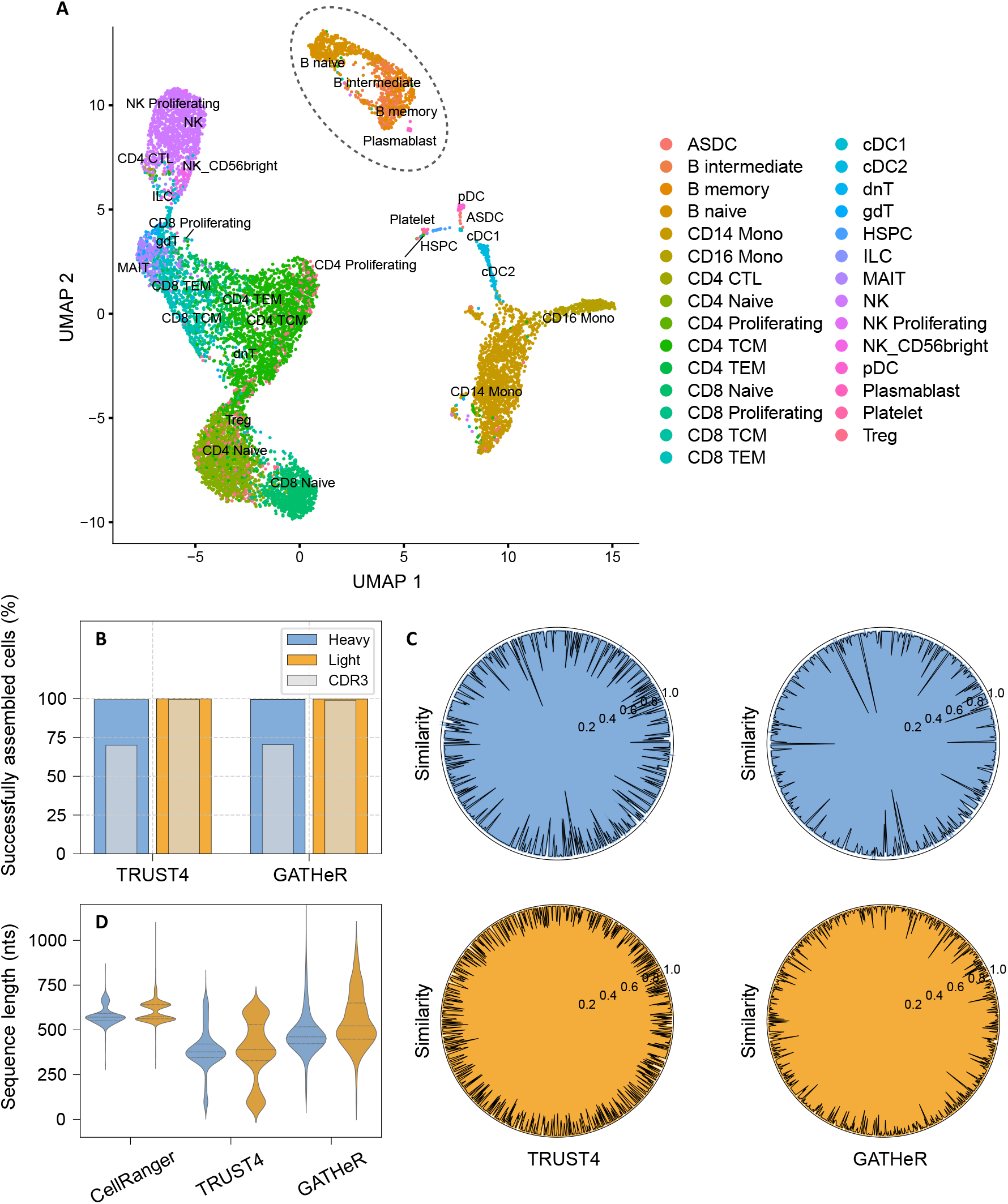
GATHeR assembles accurate BCRs from 10x Genomics 5^′^ gene expression libraries with high concordance to 10x V(D)J references. (**A**) UMAP of PBMCs generated and annotated with Seurat [18]. (**B**) Percentage of annotated B cells in which GATHeR and TRUST4 assembled heavy chains, light chains, and sequences with complete CDR3 regions. (**C**) Radar plots showing sequence similarity of heavy- and light-chain assemblies from GATHeR and TRUST4 relative to the 10x Genomics V(D)J reference. Similarity is calculated as the number of identical nucleotides divided by the length of the shorter sequence, expressed as a fraction. (**D**) Length comparison of assembled BCRs (GATHeR and TRUST4) versus the 10x Genomics V(D)J reference.

Accuracy was assessed against annotations derived from the matched 10x Genomics V(D)J-enriched library (high-confidence reference; Figure 4C). Both tools were concordant with the benchmark, with GATHeR yielding higher mean similarity (heavy: 0.97 vs. 0.95; light: 0.98 vs. 0.95). In addition, GATHeR produced longer reconstructed sequences than TRUST4 (Figure 4D), in some cases approaching lengths obtained from the pre-amplified V(D)J library.

Taken together, these results indicate that GATHeR delivers both longer and more accurate BCR reconstructions from 10x Genomics gene-expression data than TRUST4, without the need for an additional enrichment library.

## Discussion

GATHeR consistently reconstructs longer, more contiguous BCR assemblies than existing methods, with the largest gains in naive and memory B cells where Ig transcripts are sparse. By recovering the entire constant regions of both heavy and light chains, GATHeR enables reliable isotype/subclass and constant-region allele assignment, strengthens clonal lineage inference, and allows direct interrogation of membrane (M1/M2) versus secretory tailpiece usage. Extended assemblies also revealed intron retention in a measurable fraction of heavy- and light-chain transcripts that introduce premature stop codons and hence non-productivity; to our knowledge, such events have not been widely reported for heavy-chain mRNA and are overlooked by standard V(D)J callers (e.g., IgBLAST). Collectively, these results position GATHeR as a robust, end-to-end solution for sequence-resolved BCR analysis across library types and cellular states.

Although BCR diversity is often equated with V(D)J variation, human constant regions harbor substantial genetic and structural diversity, including population- and subclass-specific polymorphisms and hinge-length variants, beyond what is typically captured in repertoire studies [19, 20, 21]. Changes in heavy-chain constant domains can tune effector mechanisms by altering C1q engagement and Fc*γ*R interactions (affecting complement activation, ADCC and ADCP) [22, 23, 24], and variants that modulate FcRn binding can influence antibody transport and half-life [25]. Substitutions in CH1—and in some contexts CH2/CH3—can allosterically impact antigen binding by the Fab [26]. At the BCR itself, a cytoplasmic-tail variant (G396R) associates with altered B-cell activation, lupus risk and vaccine responses [27]. By recovering complete constant regions alongside the V(D)J, GATHeR enables reliable sub-isotype and constant-region allele calls and a direct readout of membrane versus secretory isoforms, providing a sequence-resolved framework to connect constant-region genotype with cellular function.

GATHeR revealed intron retention in Ig heavy-chain transcripts that can render them non-productive. Unproductive splicing is conserved across species [28] and has long been considered an additional layer of gene-expression control [29], yet the underlying machinery and regulatory logic remain incompletely understood. Partial intron retention was recently reported in the 5′ UTR of *κ*- and *λ*-light-chain transcripts in naive human B cells [17]. Here, we corroborate intron retention in light chains and, to our knowledge, provide the first evidence for intron retention within heavy-chain transcripts. While we cannot exclude some contribution from unspliced nuclear pre-mRNA, we also identified heavy- and light-chain transcripts with partially retained introns, consistent with bona fide splice variants rather than pre-mRNA leakage. Notably, most retained introns in heavy-chains mapped to the distal (3′) constant region—between CH3 and the membrane-coding exons—underscoring the value of full-length constant-region assembly and helping to explain why earlier V(D)J-focused surveys may have missed these events.

GATHeR assembled two or more distinct heavy- and/or light-chain transcripts in a subset of naive and memory B cells, consistent with prior single-cell reports [15]. Most events occurred in cells annotated as naive, where IgD/IgM co-expression is expected: GATHeR recovered separate productive IGHD and IGHM transcripts sharing an identical VDJ rearrangement in roughly half of naive cells, consistent with the canonical model of a single VDJ paired with alternative constant regions. Multiple chains were also observed in a minority of memory B cells. While some instances may reflect technical artifacts (e.g., doublets), the long contigs enable productivity assessment: among additional chains, 17% of heavy and 26% of light transcripts were non-productive, suggesting transcription from a second, non-productive allele that is not completely silenced.

Methodologically, GATHeR builds on a long line of assembly strategies grounded in de Bruijn graphs (DBGs). DBGs have powered short-read assembly since the Eulerian-path formulation [30] and tools like Velvet [31]; Trinity adapted DBGs to transcriptomics for isoform-aware de novo assembly [32]. In BCR reconstruction, BraCeR and BALDR apply Trinity after per-cell Ig read enrichment—preserving isoforms and SHM but at higher compute cost [9, 33]. By contrast, BASIC stitches from germline anchors, TRUST4 uses k-mer seeding with greedy extension, and MiXCR is alignment-guided [8, 10, 34]. GATHeR avoids upfront alignment/filtering and combines (i) a compacted DBG with lightweight path traversal for well-covered loci and (ii) rnaSPAdes assembly algorithm [35], which has demonstrated advantages over Trinity in producing more complete assemblies with fewer chimeras and less redundancy, while handling uneven or low coverage more effectively—capabilities that are critical for naive and memory B cells where sparse coverage and co-expression of multiple heavy chain isotypes (e.g., IgM and IgD) are common.

In conclusion, GATHeR expands what can be inferred from Ig transcripts in scRNA-seq: when constant-region reads are available, it routinely generates assemblies that span the entire constant region, detects splice-derived nonproductivity and co-expressed isoforms, and operates on both full-length and 5′ gene-expression libraries. These capabilities extend the possibilities for single-cell analyses of B cells in vaccination, autoimmunity, and malignancies.

## Methods

### Graph construction and assembly strategy

GATHeR employs compacted de Bruijn graphs (cDBGs) to facilitate the de novo assembly of RNA sequence data. The method utilizes BCALM2 [36] to compact the de Bruijn graphs (DBGs), where nodes correspond to the maximal unitigs of the original DBG. This compaction reduces cost without losing any information. Nodes are weighted based on the average abundance of the compacted *k*-mers, while edges are weighted by the mean weights of the connected nodes. The SPAdes assembler [35] is invoked when read coverage is low and/or uneven. In such cases, SPAdes-derived contigs are also incorporated into the BCR identification step.

### Error correction and graph simplification

*k*-mer errors are addressed in two steps. The first step involves the compaction process, where *k*-mers with a frequency below a certain threshold are removed. This threshold can be adjusted based on the characteristics of the data. The second step, which also contributes to graph simplification, involves removing duplicate unitigs with lower weights. Let *u* and *v* be two unitigs in a set *S*, and let *w*(*u*) and *w*(*v*) denote their respective weights. *pre*(*u, k −* 1) and *suf* (*u, k −* 1) are defined as the first and the last *k −* 1 characters of *u* respectively. Then *u* and *v* are duplicates when *pre*(*u, k −* 1) = *pre*(*v, k −* 1) or *suf* (*u, k −* 1) = *suf* (*v, k −* 1). The new set *S*\*, used to build the cDBG, contains only those *u* that have the highest weights among all satisfying the overlapping condition in *S*. Additionally, errors are implicitly corrected during the traversal and subsequent identification of the optimal path on the graph, which will be explained shortly.

### Traversing and finding optimum paths

To make traversing and pathfinding more efficient, we decompose the cDBG into a set of subgraphs by identifying weakly connected components (WCCs) using NetworkX[37]. These components are the maximal subgraphs of the cDBG where an undirected path exists between any pair of nodes. Subsequently, we assumed that each component in the set of WCCs corresponds to a single contig. Therefore, we employed the depth-first search (DFS) algorithm to traverse through the nodes and edges of each subgraph. Let subgraph *G*′ be an element of the set of WCCs. Then the path *P* comprises a sequence of nodes *P* = [*u*_1_, *u*_2_, …, *u*_*k*_] obtained from a DFS traversal of *G*′. Path *P* is considered optimal if it has the largest weighted average metric *ϕ*(*P*), which is defined as:

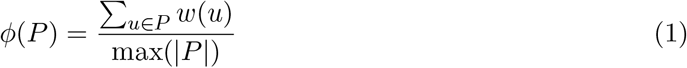

where |*P* | represents the number of nodes in the path P and max(|*P*|) is the length of the longest path in *G*.

### Immunoglobulin sequence identification

Pairwise alignment was used to map assembled sequences (contigs) to the constant and variable regions of human and mouse Ig genes, which encode BCRs. These sequences were obtained from the IMGT database. By default, the method uses the BLASTN [38] program with the following parameters for pairwise alignments: Percentage Identity (`perc_identity`) = 90%, E-value (`evalue`) *<* 0.01, and Bit Score (`bitscore`) *>* 50 Bits. GATHeR then annotates the heavy- and light-chain assemblies and outputs them, along with their corresponding weights, in FASTA format. As part of the post-processing step, GATHeR also uses a built-in aligner based on the Biopython package [39] to annotate constant domains of Igs.

### Clonality and phylogenetic analysis

As a post-processing step, an optional command uses existing tools for clonality analysis and lineage tree construction. Assembled BCR sequences from different single-cell datasets can be collected and processed using IgBLAST[40] and the Change-O[41] toolkit to comprehensively profile the Ig repertoire. This analysis identifies the most likely germline V, D, and J genes and delineates the junction regions, including the complementarity-determining region 3 (CDR3), for each sequence. Change-O uses these annotations to cluster sequences into clonotypes. Subsequently, Dowser and Igphyml are employed to construct lineage trees based on clonal assignment information combined with light chain data [42, 43].

### Processing and annotation of 10x Genomics 5^′^ gene expression and CITE-seq data

Data processing was performed in Seurat v5.2.1 [44]. A 10x Genomics feature barcode HDF5 matrix with Gene Expression and antibody capture (ADT) counts was imported; a Seurat object was created from RNA counts (min.cells = 3, min.features = 200) and ADT was added as a separate assay. Low-quality cells were removed by requiring 200–4,000 detected genes and *<* 10 mitochondrial RNA. RNA data were log-normalized (NormalizeData), variable features identified (FindVariableFeatures), scaled (ScaleData), and reduced with PCA. A shared nearest-neighbor graph was built on the first 10 PCs, clusters were called with the Louvain algorithm at resolution 0.8, and UMAP was computed on the same 10 PCs. ADT data were normalized with centered log-ratio (CLR, margin = 2). Cell types were assigned by label transfer to a multimodal PBMC reference [44] using Seurat’s SCT-based anchor mapping. Barcodes for selected clusters and predicted cell types were exported for downstream analyses.

## Data availability

All sequencing datasets analyzed in this study are publicly available. RNA-seq data for Dataset I are deposited in GEO (accession GSE116500); Dataset II in ArrayExpress (accession [E-MTAB4825](https://www.ebi.ac.uk/biostudies/ArrayExpress/studies/E-MTAB-4825)); and Dataset III in ArrayExpress (accession E-MTAB-11452). The 10x Genomics datasets are available from the [10x Genomics datasets portal](https://support.10xgenomics.com/single-cell-vdj/datasets/3.1.0/vdj_nextgem_hs_pbmc3).

## Code availability

The code is freely available at https://github.com/neuroimmunology-uio/gather.

