## Supplementary Information for "GATHeR: Graph-based Accurate Tool for Immunoglobulin HEavy- and Light-chain Reconstruction"

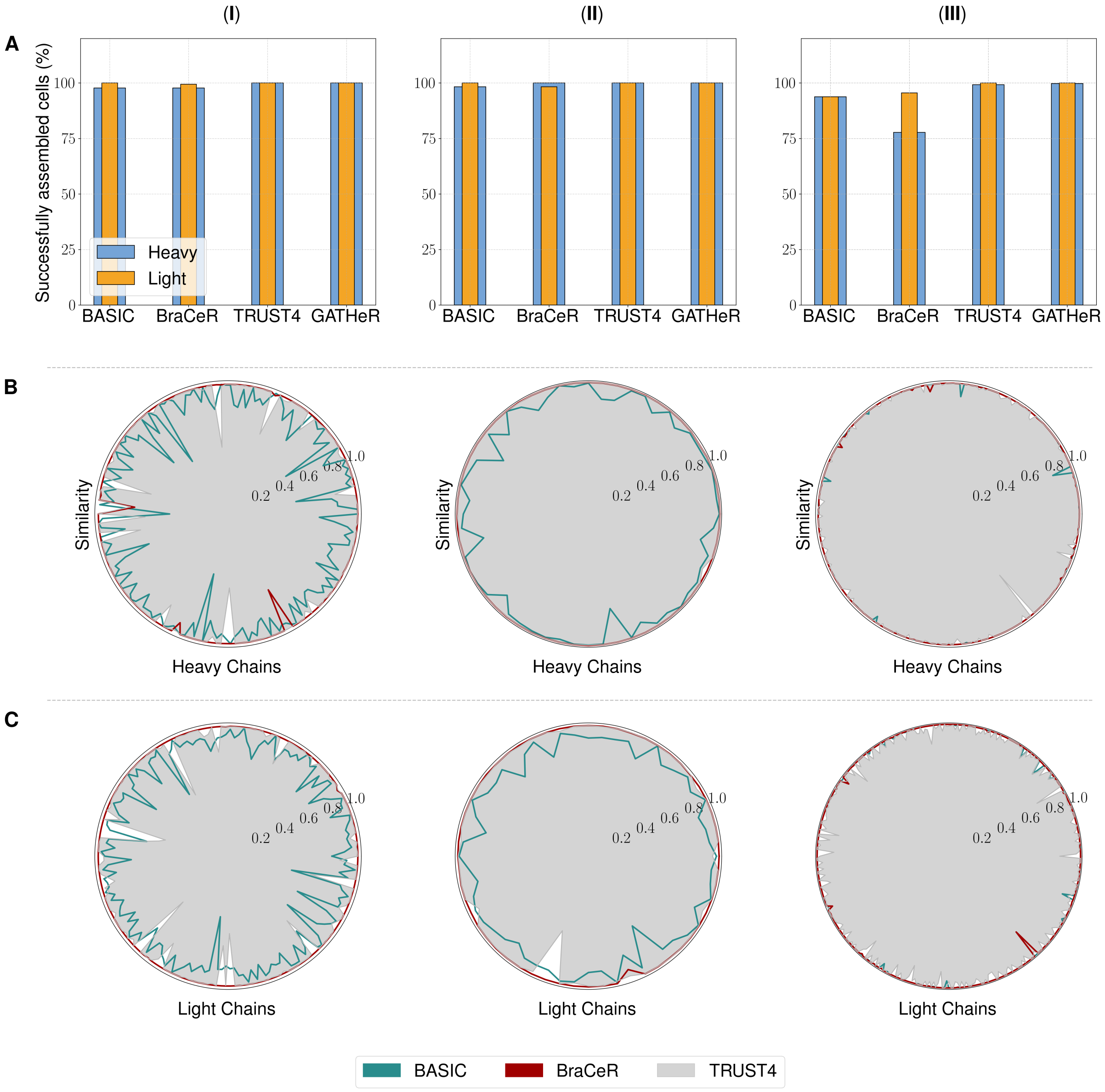


**Figure S1. Performance benchmark of GATHeR against BASIC, BraCeR and TRUST4.** (**A**) Bar plots showing, for each dataset (I–III, left to right), the fraction of cells in which each tool successfully reconstructed the heavy chain (IGH), the light chain (IGK/L) or both. (**B**) Radar plots of heavy-chain sequence similarity and (**C**) light-chain sequence similarity comparing contigs assembled by GATHeR with those from other tools across three datasets (I–III, left to right). Similarity is calculated as the number of identical nucleotides divided by the length of the shorter sequence, expressed as a fraction.

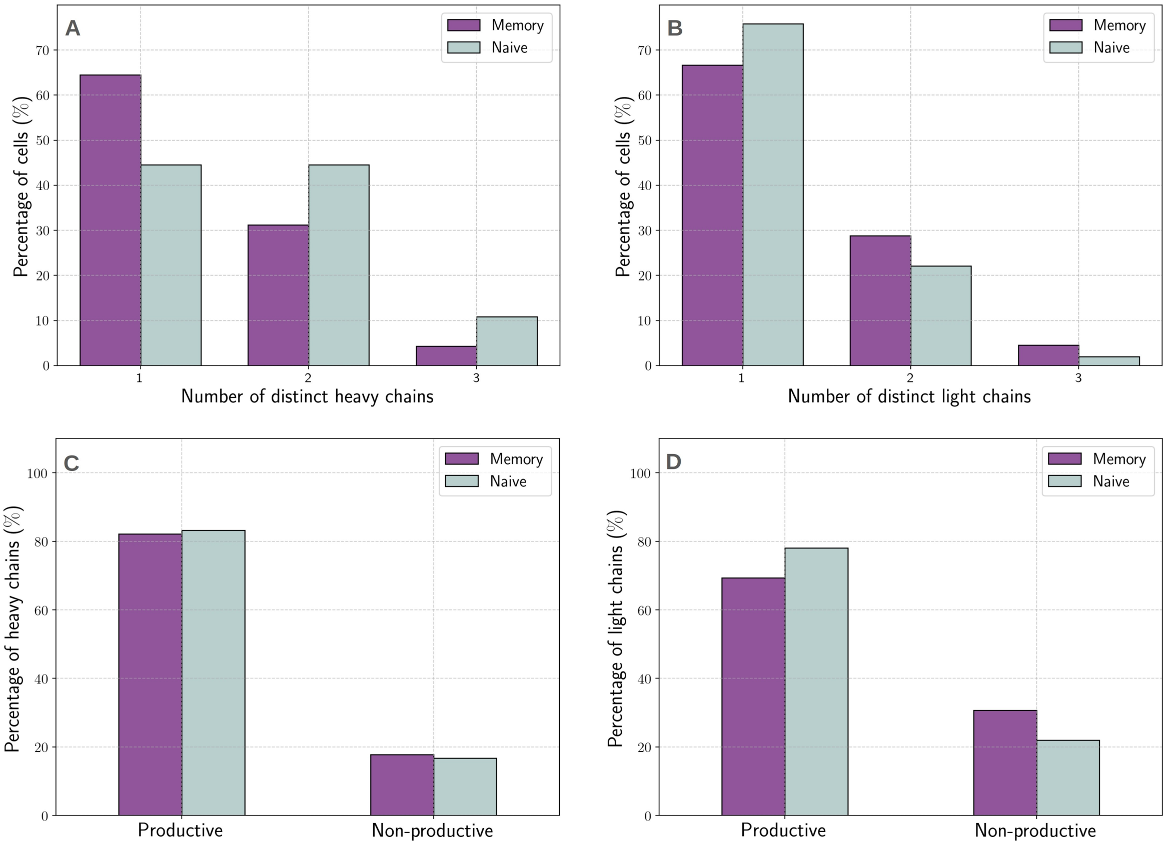


**Figure S2. GATHeR identifies multiple heavy- and light-chain sequences per cell in a subset of naive and memory B cells.** (**A**) Percentage of cells with 1, 2, or 3 reconstructed heavy-chain sequences, and (**B**) percentage with 1, 2, or 3 reconstructed light-chain sequences, shown separately for naive and memory B cells. (**C**) Percentage of productive and nonproductive heavy-chain assemblies and (**D**) light-chain assemblies in Dataset III.

### Intron retention IgM

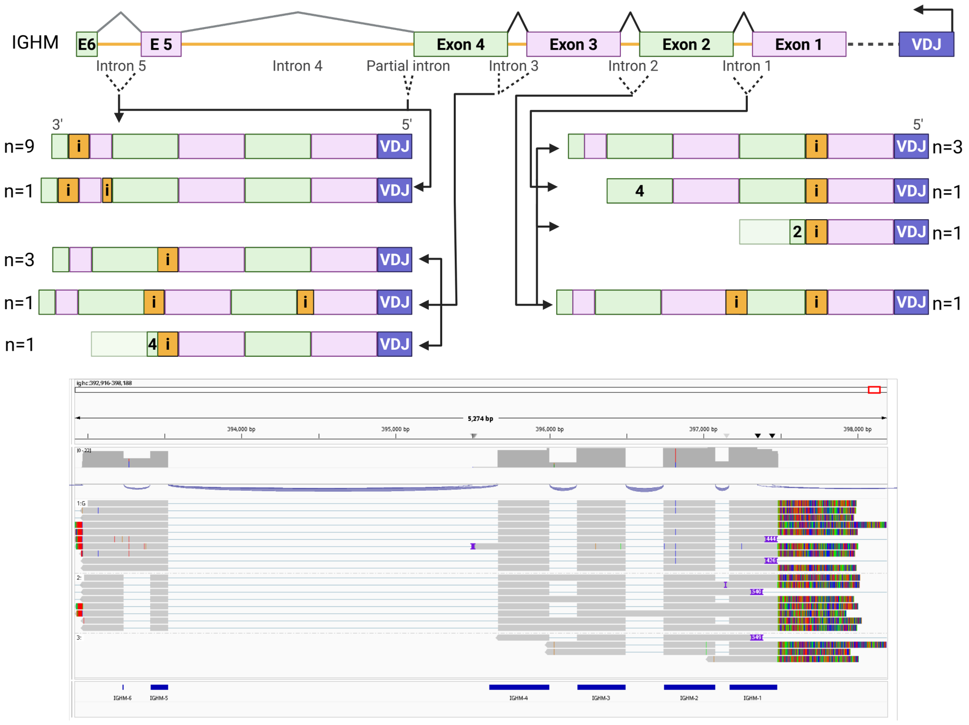


### Intron retention IgD

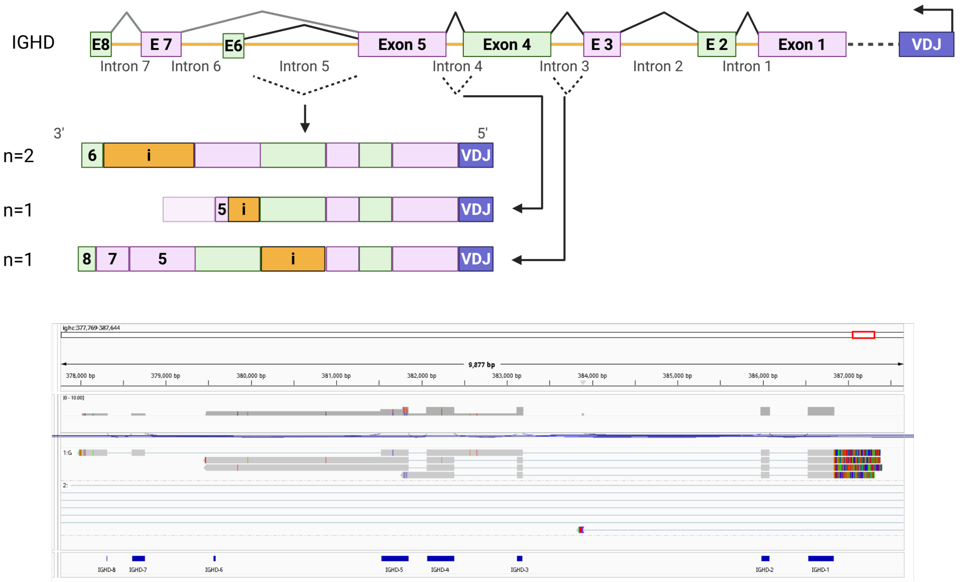


*Figure S3 (continued on next page)*

### Intron retention IgG3

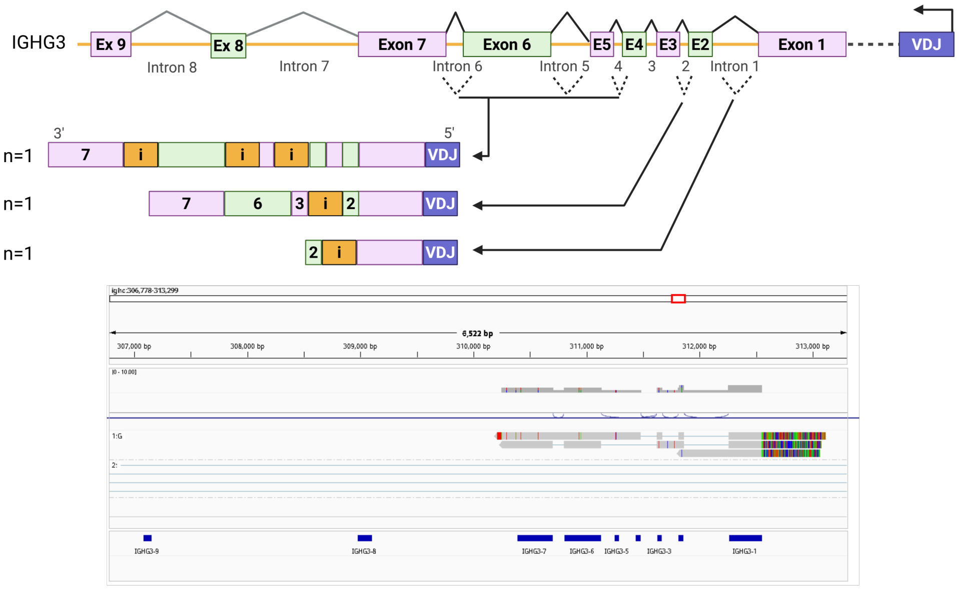


### Intron retention IgG1

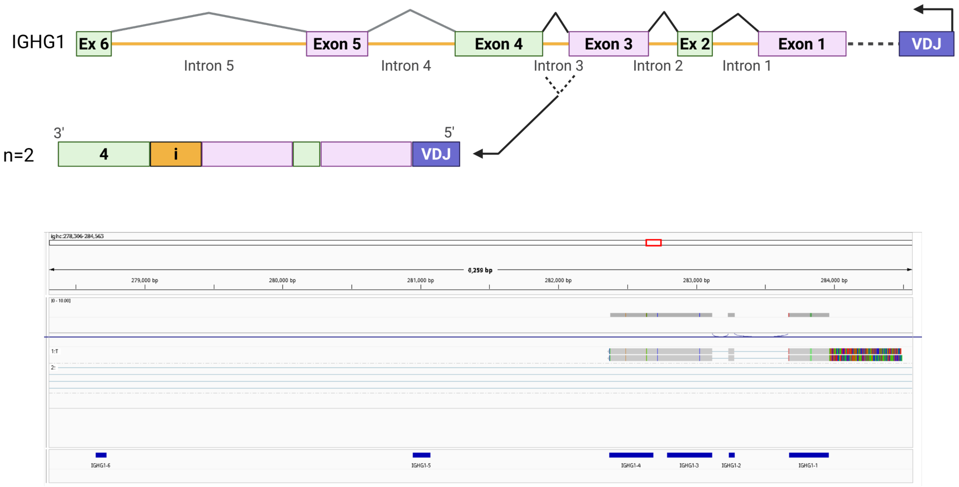


*Figure S3 (continued on next page)*

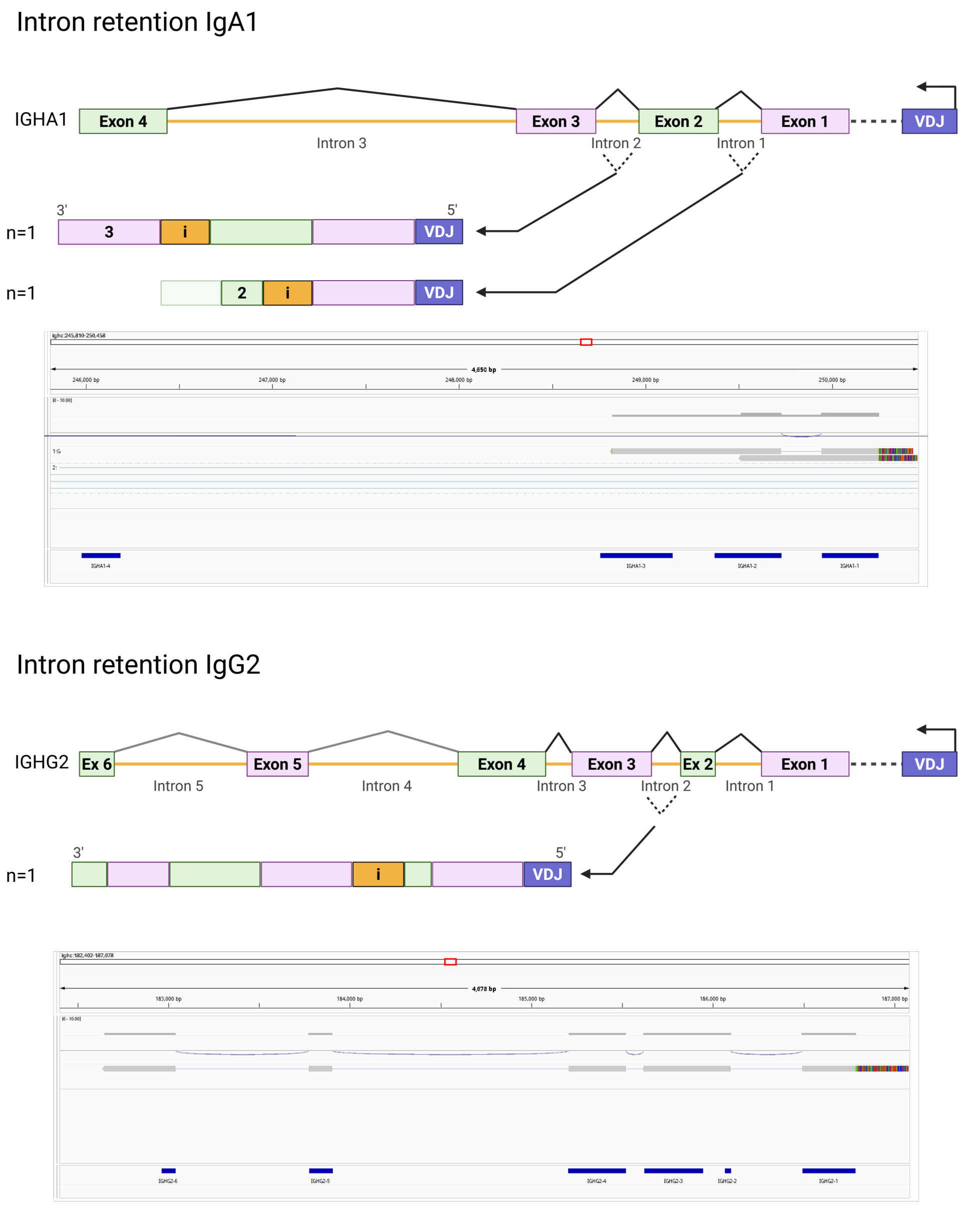


**Figure S3. Schematic of intron-retention events in GATHeR-assembled immunoglobulin heavy-chain transcripts from naive and memory B cells in Dataset III.** For each isotype (IgM, IgD, IgG3, IgG1, IgA1, IgG2), all detected splice variants are shown; numbers to the left indicate how many reconstructed variants show that splicing pattern. Below, Integrative Genomics Viewer (IGV) snapshots show how the assembled transcripts align to the corresponding germline IGH gene.

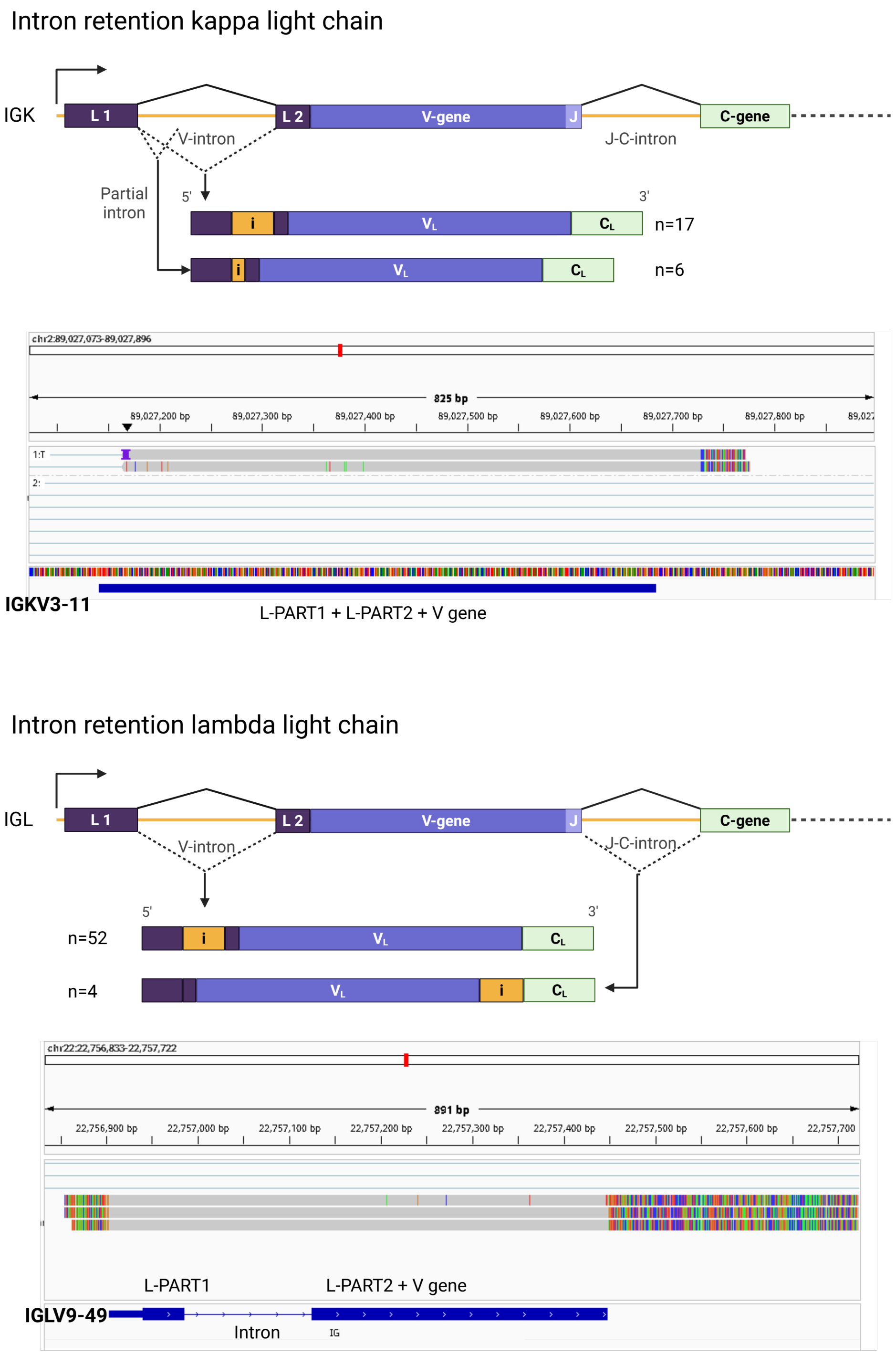


**Figure S4. Schematic of intron-retention events in GATHeR-assembled immunoglobulin light-chain transcripts from naive and memory B cells in Dataset III.** $\kappa$ chains exhibit partial and complete L–V (leader–variable) intron retention, whereas $\lambda$ chains show L–V and J–C (joining–constant) intron retention. Numbers to the left indicate how many reconstructed variants show that splicing pattern. Integrative Genomics Viewer (IGV) snapshots below illustrate alignments of the assembled transcripts to the corresponding germline *IGK*/*IGL* genes.

| donor | sequence_id | c_call | CH2_idn | CH4_idn | CH3_idn | CH1_idn | M1_idn |
|---|---|---|---|---|---|---|---|
| donor1 | cell_1 | IGHM*03-N | 0.99 | 1.0 | 1.0 | 1.0 | 1.0 |
| donor1 | cell_2 | IGHM*03-N | 0.99 | 1.0 | 1.0 | 1.0 | 1.0 |
| donor1 | cell_3 | IGHM*03-N | 0.99 | 1.0 | 1.0 | 1.0 | 1.0 |
| donor1 | cell_4 | IGHM*03 | 1.00 | 1.0 | 1.0 | 1.0 | 1.0 |
| donor1 | cell_5 | IGHM*03-N | 0.99 | 1.0 | 1.0 | 1.0 | 1.0 |
| donor1 | cell_6 | IGHM*03 | 1.00 | 1.0 | 1.0 | 1.0 | 1.0 |
| donor1 | cell_7 | IGHM*03-N | 0.99 | 1.0 | 1.0 | 1.0 | 1.0 |
| donor2 | cell_1 | IGHM*03 | 1.00 | 1.0 | 1.0 | 1.0 | 1.0 |
| donor2 | cell_2 | IGHM*03 | 1.00 | 1.0 | 1.0 | 1.0 | 1.0 |
| donor2 | cell_3 | IGHM*03 | 1.00 | 1.0 | 1.0 | 1.0 | 1.0 |
| donor2 | cell_4 | IGHM*03 | 1.00 | 1.0 | 1.0 | 1.0 | 1.0 |
| donor2 | cell_5 | IGHM*03 | 1.00 | 1.0 | 1.0 | 1.0 | 1.0 |
| donor2 | cell_6 | IGHM*03 | 1.00 | 1.0 | 1.0 | 1.0 | 1.0 |
| donor2 | cell_7 | IGHM*03 | 1.00 | 1.0 | 1.0 | 1.0 | 1.0 |

**Table S1. Simplified output table from GATHeR analysis of constant regions in assembled heavy chains from 14 naive B cells derived from two donors in Dataset III.** The novel allele in donor 1 is annotated with the suffix "-N". Columns `CH2_idn`, `CH4_idn`, `CH3_idn`, `CH1_idn`, and `M1_idn` indicate the sequence identity of the assembled regions relative to the IMGT database.
